# Visual short-term memory related EEG components in a virtual reality setup

**DOI:** 10.1101/2023.01.23.525140

**Authors:** Felix Klotzsche, Michael Gaebler, Arno Villringer, Werner Sommer, Vadim Nikulin, Sven Ohl

## Abstract

Virtual reality (VR) offers a powerful tool for investigating cognitive processes, as it allows researchers to gauge behaviors and mental states in complex, yet highly controlled, scenarios. The use of VR head-mounted displays in combination with physiological measures such as EEG presents new challenges and raises the question whether established findings also generalize to a VR setup. Here, we used a VR headset to assess the spatial constraints underlying two well-established EEG correlates of visual short-term memory: the amplitude of the contralateral delay activity (CDA) and the lateralization of induced alpha power during memory retention. We tested observers’ visual memory in a delayed match-to-sample task with bilateral stimulus arrays of either two or four items while varying the horizontal eccentricity of the memory arrays (4, 9, or 14 degrees of visual angle). The CDA amplitude differed between high and low memory load at the two smaller eccentricities, but not at the largest eccentricity. Neither memory load nor eccentricity significantly influenced the observed alpha lateralization. We further fitted time-resolved spatial filters to decode memory load from the event-related potential as well as from its time-frequency decomposition. Classification performance during the retention interval was above chance level for both approaches and did not vary significantly across eccentricities. We conclude that commercial VR hardware can be utilized to study the CDA and lateralized alpha power, and we provide caveats for future studies targeting these EEG markers of visual memory in a VR setup.

**Impact statement:** Combining EEG with virtual reality, we studied how the eccentricity of a memory array during encoding affects well-known neural markers of visual short-term memory. We reveal that the previously reported occurrence of these EEG components during visual memory retention can be replicated in such a setup. These EEG markers were differently affected by eccentricity, hence providing valuable constraints for future experimental designs.

## Introduction

In every moment, we are surrounded by a wealth of visual signals that can guide our behavior, regardless of whether they are continuously present, temporarily occluded, or have just disappeared from view. For instance, when waiting at a crosswalk, we visually assess the distance to the opposite side of the street, the color that the traffic light is currently displaying, and whether cars are approaching from either side. Now, if a car suddenly passes us, it will occlude relevant parts of the scene and we can no longer directly access these previously available visual signals. Such information, however, is available in visual short-term memory which allows us to maintain representations of the most relevant objects in the visual scene (e.g., the number of approaching cars) and use them for informing our actions. Several fundamental functions of visual cognition rely on visual short-term memory. As an example, consider the challenge to compare the visual impressions of the street scene before and after the car blocked our view. The capacity of visual short-term memory, however, is limited, which implies that only a small fraction of the visual details in our environment can be maintained (Cowan, 2010; Luck & Vogel, 1997; Marois & Ivanoff, 2005; Pashler, 1988).

A powerful window into this domain of human cognition is provided by neuroscientific approaches, such as electroencephalography (EEG), that offer physiological markers for visual short-term memory processes. One of the best-established markers is the *contralateral delay activity* (CDA), a lateralized component of the event-related potential (ERP). Its amplitude increases with the number of items maintained in visual short-term memory and reflects the individual short-term memory capacity (Luria et al., 2016; Vogel & Machizawa, 2004). In addition to the CDA, the power of oscillations in the alpha frequency range (8–13 Hz) constitutes a further component of the EEG signal that has been associated with visual short-term memory (Pavlov & Kotchoubey, 2022; Sauseng et al., 2009; Woodman et al., 2021). More specifically, after encoding lateralized stimuli, alpha oscillations recorded in EEG channels contralateral to the memorized stimulus exhibit less power than those in ipsilateral channels (de Vries et al., 2020; Hakim et al., 2019; Leenders et al., 2018). The relationship between this lateralization of alpha power, the CDA, and the aspects of visual short-term memory that they each reflect has been discussed several times (Fukuda et al., 2015; Hakim et al., 2019; Nikulin et al., 2007; van Dijk et al., 2010), but has not yet been conclusively unraveled.

Using immersive virtual reality (VR) promises to approximate more naturalistic (i.e., less constrained) contexts for the investigation of the CDA and lateralized alpha oscillations as well as their role for visual short-term memory than standard laboratory experiments. VR technologies offer the possibility to simulate complex and interactive spatial scenarios under rigorous experimental control in laboratory conditions. By tracking body movements (e.g., of head and limbs), a VR setup can dynamically adapt the computer-generated stimuli, thereby modelling the spatial structures of the surroundings and the constant interaction with them. Current VR headsets use two screens—one for each eye—to display virtual content, enabling stereoscopic depth perception. In addition, VR allows researchers to easily collect various motion measures (e.g., head and hand movements, gaze patterns) and to link them to the virtual scene as well as to synchronously collected neurophysiological data streams (e.g., EEG). For example, combining eye tracking and VR allowed to disentangle the contributions of retinotopic and spatiotopic memory representations around self-movements (Draschkow et al., 2022b), demonstrating how experiments in the context of natural behavior can inform studies of visual short-term memory (see Kristjánsson & Draschkow, 2021, for review). However, testing visual short-term memory with a VR headset while simultaneously recording EEG poses several challenges for the EEG setup (Cattan et al., 2018; Tauscher et al., 2019; Weber et al., 2021). For instance, the headset exerts a mechanical effect on the physiological sensors, emits electromagnetic radiation, imposes a weight on the wearer that leads to tensing of head and neck muscles, and invites the observer to explore the surrounding virtual environment which is usually accompanied by (head, eye, and body) movements. All of these aspects are known to impact the signal quality of EEG (Luck, 2014). It is therefore unclear whether the EEG components related to visual short-term memory processing that can be observed in a conventional lab setting are also observable when using a VR headset.

Furthermore, studies investigating EEG correlates of visual short-term memory have typically presented the stimuli either in foveal or parafoveal vision. It is therefore not clear how EEG components like the CDA or alpha power lateralization depend on stimulus eccentricity and if they occur in a memory task that includes larger eccentricities. A better understanding of such spatial constraints is particularly interesting for VR studies, given that immersive setups aim to evoke the impression of being surrounded by the virtual world through stimulating the largest possible part of the visual field. Especially in setups which allow the participant to move, relevant stimuli may therefore occur at various retinotopic locations, well into the periphery of visual perception. With larger receptive fields in the periphery as compared to foveal and parafoveal vision (Freeman & Simoncelli, 2011), peripheral stimuli are processed by fewer neurons and could thereby evoke altered neurophysiological responses. Also, stimuli at different eccentricities will elicit responses in different parts of the retinotopically organized visual cortices, which might, due to cortical folding, result in altered amplitudes or topographies of EEG responses. Indeed, various EEG components vary with stimulus eccentricity (Bahramisharif et al., 2011; Busch et al., 2004; Capilla et al., 2016; Domínguez-Martínez et al., 2015; Papaioannou & Luck, 2020). For example, in a recent experiment studying visual attention, Papaioannou & Luck (2020) showed that the *Post-N2pc Positivity* (PNP), a lateralized component of the ERP, exhibits higher amplitudes for stimuli with larger eccentricities. However, whether and how also EEG components related to visual short-term memory vary as a function of stimulus eccentricity has not yet been investigated. A strong modulation by eccentricity or even their absence beyond a particular eccentricity would constitute a major constraint for future studies combining EEG and VR in visual short-term memory tasks.

In the present study, we asked whether the CDA and lateralized alpha power—two prominent EEG markers of visual short-term memory—can be observed in a well-controlled delayed match-to-sample task implemented in a VR setup and whether they depend on the eccentricity of the memory stimulus. Classical inferential analyses and recently introduced multivariate decoding methods revealed the presence of the CDA, a lateralization of alpha power, and the availability of information about the memory load in lateralized and non-lateralized features of the EEG signal. Hence, despite the technical challenges of neurophysiological recordings in a VR setup, visual short-term memory related EEG components can still be observed. However, the results were less conclusive for stimuli presented at an eccentricity of 14 dva compared to when they were presented at eccentricities of 4 and 9 dva, suggesting that large eccentricities have an impact on some of the components. Overall, we provide a basis for future studies of visual short-term memory using EEG together with VR hardware. All data and code necessary to reproduce our results are publicly available: https://osf.io/btrws/.

## Method

### Participants

We tested 26 naïve participants (20–38 years, *M* = 26.62, *SD* = 4.92, 12 female, 21 right-handed) with normal or corrected-to-normal vision according to self-report. Persons wearing glasses could not participate in the study due to incompatibility with the eye tracker. A total of 21 out of these 26 participants contributed to the final sample: Two participants opted out before the end of the experiment, and we excluded the data of three participants based on an excessive number of rejected trials (as described in the section *EEG preprocessing*).

Using Ishihara’s color test (Clark, 1924) and a Titmus test (*Fly-S Stereo Acuity Test*, Vision Assessment Corporation, Hamburg, Germany), we ensured participants’ intact color vision and stereopsis. None of the participants reported psychiatric, neurological, or cardio-vascular conditions. The study was approved by the ethics committee of the Department of Psychology at the Humboldt-Universität zu Berlin and participants provided their written consent prior to participation. Participants were compensated with 9€ per hour. The experimental session took 4.5 hours on average.

### Materials

The experiment was conducted using a VR head-mounted display (HTC Vive Pro, HTC, Taiwan) with one AMOLED display per eye (refresh rate: 90 Hz, spatial resolution: 1440 x 1600 px, field of view: 55 degrees of visual angle; for an in-depth discussion of relevant hardware parameters see Lynn et al., 2020). At the beginning of the session, the VR headset was adjusted to match the interpupillary distance of each individual participant. We tracked participants’ eyes with a sampling frequency of 200 Hz using the Pupil Labs VR/AR eye tracking add-on mounted within the headset (Pupil Labs, Berlin, Germany). Participants reported their memory by pressing one of two buttons (the “trigger button” or the “trackpad”) on one handheld controller of the HTC Vive Pro VR system. The assignment of the function of the two buttons, as well as the choice of whether to use the left or the right hand, was randomized and counterbalanced across participants.

For the implementation of the experiment, we used the Unity software (v2018.3.11; Unity Technologies, San Francisco, United States) in combination with the SteamVR Unity plugin (v2.0.1, Valve Corporation, Bellevue, United States) and ran it on a VR-ready PC (Intel Core i9-9900K, 3.6 GHz, 32 GB RAM, NVIDIA RTX 2080Ti GPU, Windows 10). The computer was connected to the EEG amplifier via an analogue port (D-SUB 25) to enable synchronization of the EEG data with the experimental events. We used the Unity Experiment Framework (UXF; Brookes, 2017/2019; Brookes et al., 2020) for structuring the experiment and recording events as well as for tracking the behavioral data (i.e., memory responses, head and controller movements). The eye tracker was interfaced from the experimental code using the Unity plugin hmd-eyes (Pupil Labs, 2016/2019) and the Pupil software (Pupil Labs, 2013/2019; Kassner et al., 2014). To synchronize the data streams (i.e., behavioral reports, eye positions, EEG), we used custom C# scripts and network-based communication (i.e., timestamps in the eye tracking data) as well as analogue triggers (EEG). We recorded EEG and electrooculogram (EOG) data using the BrainVision Recorder software (v1.22.0101; BrainProducts, Gilching, Germany). Before and after the VR-based part of the experiment, participants filled in questionnaires implemented in SoSciSurvey (Leiner, 2019).

Participants’ EEG was sampled at a rate of 500 Hz (including a hardware-based lowpass filter at 131 Hz; third order sinc filter, –3 dB cutoff) with a 64-channel LiveAmp and actiCAP snap electrodes (both by BrainProducts, Gilching, Germany) from 60 scalp locations according to the International 10/20 system. We further measured the horizontal and vertical EOG with four electrodes attached next to the outer canthi and below both eyes. All electrodes were referenced to electrode FCz (ground: FPz). At the beginning of each experimental session all impedances were below 25 kΩ. The VR headset was carefully placed on top of the EEG cap which was covered by a disposable shower cap. The flat design of the actiCAP snap electrodes and the large cushion on the HTC Vive Pro covering the back of the head allowed for a broad, evenly spread weight distribution without excessive pressure on any single electrode. A customized facial interface cushion (VR Cover) with recesses at the appropriate sites helped to avoid pressure on the frontal EEG electrodes (Fp1/2) by the facial mask of the headset.

Due to the in-built Fresnel lenses in the VR headset and the prewarping of the image to counteract distortions by the lenses, the size and position of stimuli in terms of dva cannot simply be calculated from their size and position expressed in pixels on the displays. Therefore, we used the units of the virtual environment for calculating size and positions of the stimuli. Distances in the virtual environment are approximately equivalent to distances in the physical environment and the hardware setup of the HTC Vive Pro allows for a minimal angular resolution of 0.041 dva (Lynn et al., 2020). *Stimuli*. We presented all stimuli against a uniform grey background on an imaginary sphere (diameter: 1 m) surrounding the head of the participant in the virtual space. The white fixation symbol was a combination of a cross-hair (diameter: 1 dva) and a bull’s eye (Thaler et al., 2013) which we presented in the center of the field of view of the headset (irrespective of the position and rotation of the participant’s head). The memory and distractor items were colored circles with a diameter of 0.8 dva. In each trial, the colors were randomly chosen (without replacement) from a set of nine perceptually distinct colors as defined in CIE-LAB color space (Hinshaw, 2012; Holy, 2011). To determine the positions of the items in the memory array, we randomly selected horizontal and vertical coordinates within a spherical rectangle (with a width of 4 dva and a height of 8 dva) and ensured that the center points of adjacent circles were at least 1.6 dva apart.

### Procedure

We tested participants’ visual working memory in a delayed match-to-sample task with bilateral stimulus arrays. More specifically, participants memorized the colors of two or four circles presented in the cued visual hemifield and reported, after a retention interval, whether one of the colors had changed in a subsequent memory test. At the beginning of each trial, we presented participants with the central fixation symbol and instructed them to maintain fixation on this symbol throughout the duration of the trial (**Figure 1a**). After an initial fixation period of 800 ms, the fixation symbol turned into an arrow cue for 800 ms pointing with equal probability either to the left or to the right. Subsequently, either two or four colored circles (i.e., low vs high memory load, respectively) appeared in each visual hemifield for 200 ms. Participants were instructed to remember only the circles displayed in the hemifield initially cued by the arrow (i.e., the memory array) and to ignore the ones in the opposite hemifield (i.e., the distractor array). The arrays appeared at an eccentricity of either 4, 9, or 14 dva—as defined for the center of the invisible rectangles—measured along the horizontal meridian and relative to the fixation cross. Therefore, they had fixed coordinates relative to the participant’s field of view, independent of changes in head location or orientation (i.e., head-contingent presentation). Stimulus eccentricity was manipulated orthogonally to memory load. Following 2,000 ms after stimulus offset, we probed participants’ memory by displaying the memory arrays again with the same spatial layout. Importantly, in half of the trials, one of the circles in the memory array (i.e., in the cued hemifield) showed a different color which was randomly chosen from the remaining set of colors not yet used in this trial. Participants reported in a yes-no format whether any of the circles in the cued hemifield had changed its color.

**Figure 1.**
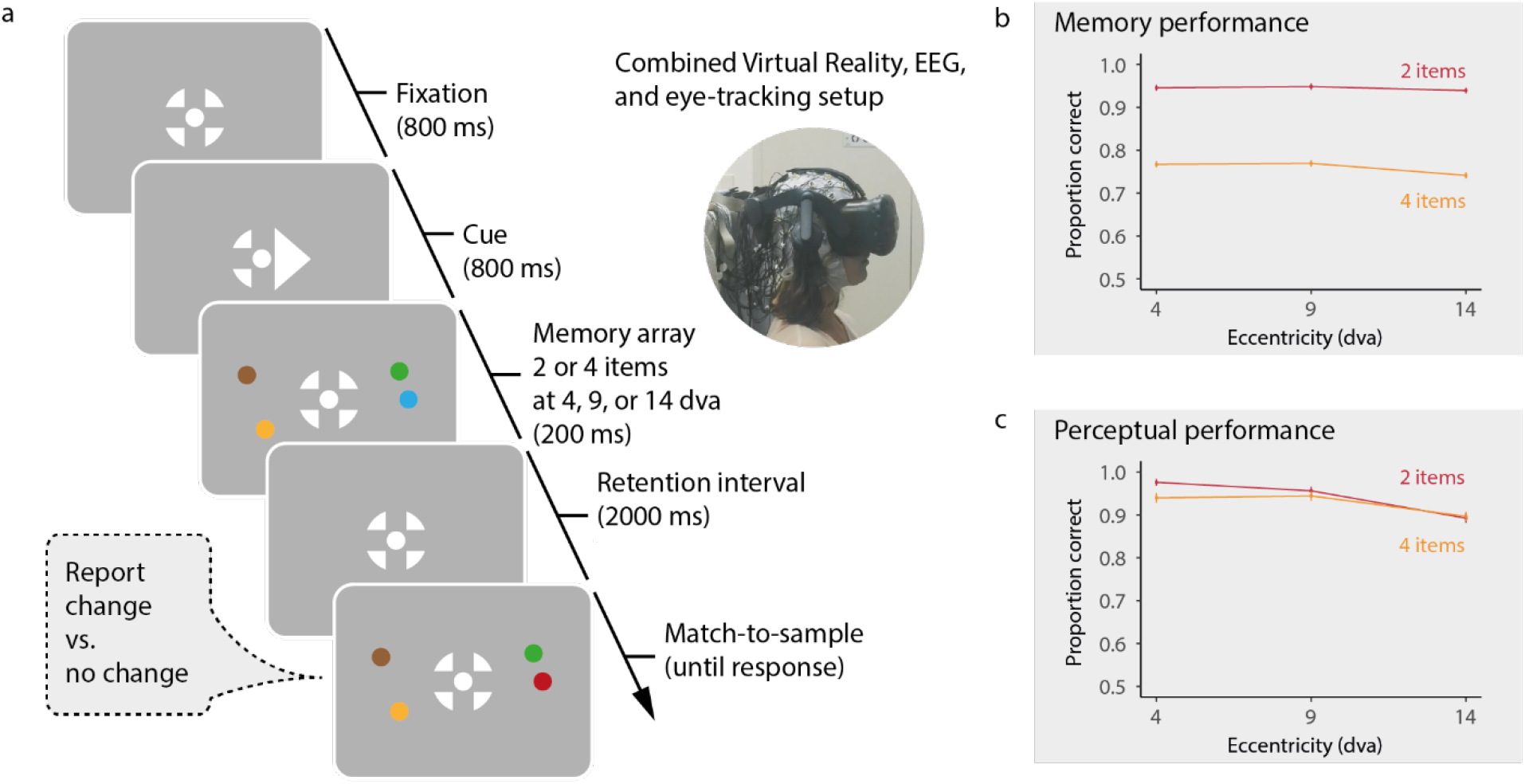
(a) The design of a single trial with low memory load in the delayed match-to-sample task and the experimental setup. For the perceptual control trials, the retention interval was omitted. (b) Behavioral results of the memory task and (b) the perceptual task. Error bars in (b) indicate ±1 SEM, taking into account the repeated measures design (Baguley, 2012; Morey, 2008).

At the beginning of the experimental session, 10 training trials allowed the participants to familiarize themselves with the task. The main experiment consisted of 10 blocks with 72 trials each. Between blocks, participants could take off the VR headset and rest until they felt ready to continue. We tested the two memory load conditions and the three stimulus eccentricities in randomly interleaved trials. Each combination of memory load and stimulus eccentricity occurred 12 times per block, resulting in a total of 720 trials in the main memory task. At the beginning of each block and after every 24 trials, we calibrated the eye tracker. In case of poor eye tracking quality, the experimenter launched additional calibrations. Based on the recommendations by Pupil Labs, 16 calibration targets were sequentially displayed in the center of the field of view and on three concentric circles (radii: 2, 4, and 8 dva) surrounding it. Noteworthy, the real-time headtracking of the VR system allowed us to present all stimuli in relation to the current position of the participant’s head. Hence, also small head movements during a trial did not displace the stimuli within the participant’s field of view.

Prior to the main experiment, participants performed the same task but without any retention interval. In half of these trials, immediately after displaying the memory array for 200 ms, one of the circles on the cued side changed its color. This change detection task tested the perceptual difficulty of the paradigm in the VR setup and whether stimuli and potential color changes were equally recognizable at all eccentricities. We did not record EEG in these perceptual control trials. Again, participants performed 10 training trials to get acquainted with the task, before running 72 trials of the perceptual task.

## Data analysis

### Behavioral data

All analyses of behavioral data were completed using the R environment (R Core Team, 2021) and RStudio (RStudio Team, 2021). To assess the effects of memory load and stimulus eccentricity on memory performance, we performed a repeated measures analysis of variance (rmANOVA) with the within-participant factors memory load (low: 2 items, high: 4 items) and stimulus eccentricity (4, 9, 14 dva). These analyses were conducted separately for data from the memory task and the perceptual task. To further assess significant effects in the ANOVA, we calculated pairwise, two-tailed *t*-tests between the single experimental conditions.

### Eye tracking data

We analyzed eye positions in an interval from 200 ms before to 2,200 ms after memory array onset using custom scripts in R. To account for the spherical presentation of the stimuli, the gaze vectors for each eye were translated to spherical coordinates (inclination, ρ, and azimuth, θ) with the origin set to the center of the respective eyeball (reconstructed by the 3D pupil detection during recording; Pupil Labs, 2013/2019) and the zenith (ρ = θ = 0 dva) set to the position of the fixation cross. Here, horizontal and vertical gaze deviations from the fixation cross correspond to azimuth and elevation, respectively, of the gaze vectors per eye. Before detecting saccades, we linearly interpolated eye tracking samples with confidence values—provided by the Pupil Labs software—smaller than 0.60 (range: 0–1.00). Whenever more than 100 consecutive samples were affected by low confidence, they were not interpolated. Saccade detection was based on a velocity-based algorithm with noise-dependent threshold (i.e., 6 SDs velocity threshold and minimum duration of 4 samples; Engbert et al., 2015; Engbert & Kliegl, 2003; Engbert & Mergenthaler, 2006). We detected saccades using only data from the eye with higher average confidence values throughout the trial and ignored saccades that overlapped with blinks (i.e., 100 ms before and after the blink). Blinks were identified by using the confidence-based blink detection implemented in the Pupil Player software (Pupil Labs, 2013/2019). We rejected any trials from all further statistical analyses (but not from EEG preprocessing), which contained a saccade with an amplitude of 2 dva or larger.

### EEG preprocessing

For the EEG data analyses, we used MNE-Python (v0.24; Gramfort et al., 2013), scikit-learn (v1.0.2; Pedregosa et al., 2011), and NumPy (v1.21.5; Harris et al., 2020). Eye movement and blink artifacts in the EEG recordings were removed using ICA decomposition (extended Infomax). To improve the fit of the ICA, we made a copy of the raw data and filtered it between 1 and 40 Hz (FIR filter with a hamming window of length 1651 samples, lower/upper passband edge: 1.00/40.00 Hz, lower/upper transition bandwidth: 1.00/10.00 Hz, lower/upper −6 dB cutoff frequency: 0.50/45.00 Hz). From this copy we removed epochs with particularly noisy EEG signals which we identified using the *autoreject* software (v0.3; Jas et al., 2017). No data interpolation was performed in this step and the rejected EEG epochs were only discarded from the copy of the data used for determining the ICA decomposition. After fitting the ICA, we identified and removed the two components with the highest correlation with the bipolar EOG channels. The remaining ICA weights were used to clean a separate copy of the continuous data which we filtered between 0.1 and 40 Hz (FIR filter with hamming window of length 16501 samples, lower/upper passband edge: 0.10/40.00 Hz, lower/upper transition bandwidth: 0.10/10.00 Hz, lower/upper −6 dB cutoff frequency: 0.05/45.00 Hz). For the CDA analysis, we extracted epochs of 2,900 ms length from −600 to 2,300 ms, relative to the onset of the memory array. For the analysis of lateralized alpha power, we extracted longer epochs, from −1,400 to 2,500 ms relative to stimulus onset, entailing the cue period as well as some buffer on both ends to reflect signal changes already in response to the cue and to avoid edge artifacts when performing wavelet convolution. We baseline-corrected these data by subtracting the mean voltage during the baseline intervals (i.e., 200 ms before stimulus/cue onset for CDA/alpha power analysis, respectively) and used autoreject for local (i.e., per participant, sensor, and epoch) interpolation and trial rejection. Finally, all trials with bad EEG or the presence of saccadic eye movements were removed. Importantly, EEG- and eye tracking-based rejections were performed independently, allowing also trials with saccadic eye movements to inform ICA decomposition. We excluded participants with more than 20% of rejected trials (n = 3) from all further analyses. For the remaining participants, 1,736 trials were rejected due to eye movements or poor EEG quality, leaving 13,384 trials in total for further EEG analysis (*M* = 88.52% per participant, range: 73.47–98.61%).

### CDA analysis

For the CDA analysis, we pooled the data from the two cuing conditions (i.e., memory array in the left or right hemifield) and computed CDA amplitudes by specifying a bilateral region of interest (ROI) that comprised parietal and occipital electrodes (P3/4, P5/6, PO3/4, PO7/8, O1/2). The selection of this ROI followed Hakim et al. (2019) who investigated the same EEG components collected with a conventional lab setup. On a single trial basis, we calculated the lateralized signal by subtracting the average signal in channels ipsilateral to the memory stimulus from the average signal in contralateral channels. The temporal evolution of the CDA was assessed using the *collapsed localizer* approach suggested by Luck & Gaspelin (2017). To this end, we pooled the data across all experimental conditions (all memory loads and eccentricities) and applied cluster-based permutation testing to identify time windows of strong lateralization. We calculated the average CDA wave form for each participant and, on the group level, applied two-tailed, one-sample *t*-tests against zero for each time point during the retention interval. Adjacent time points with *p*-values smaller than the initial threshold (0.05) were summarized as a cluster by adding their *t*-values. Significance of the clusters was determined by repeated *sign flipping*, that is in 10,000 permutations, we multiplied each of the CDA time series (averaged per participant and cropped to the retention time window) randomly either with 1 or −1. The mass of the largest cluster in each repetition was used to form a null distribution (i.e., for the hypothesis of no difference between contra- and ipsilateral signals). For statistical significance testing, we compared the cluster masses computed for the unshuffled data with this distribution. This allowed us to test whether the grand-average CDA amplitude (i.e., across all experimental conditions) was significantly different from zero (*p* < .05; cluster-corrected) during the retention interval and which time windows were driving this effect. To assess the effects of the experimental manipulations (memory load, stimulus eccentricity) and their interaction, we calculated the mean CDA amplitude in the identified time windows and modeled it on the participant-level with a rmANOVA.

### Decoding from the ERP

We complemented the univariate statistics by a multivariate approach to increase the sensitivity of the analyses. By fitting a sliding linear classifier, we decoded at various time points during the retention interval whether the sample was measured during the maintenance of a high or low memory load (Adam et al. 2020). The samples consisted of the average ERP in mini-batches of 10 trials from the same memory load condition. Using averaged data from mini-batches has been found to improve the performance of classifiers trained on EEG data (Adam et al., 2020; Grootswagers et al., 2017) due to suppression of noise by averaging. Furthermore, the signal was downsampled by calculating the average voltage in subsequent time windows of 10 samples (i.e., 20 ms). For each time window, we then trained a logistic regression model (solver: liblinear, L2-regularization: *λ* = 1.0) on the data from all 60 EEG sensors. The decoding performance was assessed using a randomized 5-fold cross-validation (CV) regime. That is, in each of 5 iterations, we trained the model on 80% of the averaged mini-batches and assessed its predictive performance on the remaining 20%. The area under the curve of the receiver operating characteristic (ROC-AUC), averaged across CV folds, served as a measure for the decoding performance. We repeated this decoding procedure 100 times for each participant and time window to obtain a more robust estimate. Random allocation of trials into mini-batches and random split of mini-batches into test and training sets were performed in each repetition. An overall decoding score per time window and participant was obtained by averaging the ROC-AUC scores across all repetitions. For a physiological interpretation of the decoding results, we calculated the “patterns” of the classifier (by multiplying the covariance of the EEG data with the filter weights; Haufe et al., 2014) for each time window, repetition, and participant. To determine if the decoding performance was significantly higher than chance level during the retention interval, a cluster-based permutation test was conducted on the group level using the same procedure as outlined for the CDA analysis. For each time window, we assessed whether the decoding scores per participant were significantly higher than chance level by conducting one-sided paired *t*-tests. To obtain an empirical estimate of the chance level of the classifier (for each participant and time window), the same classification procedure was conducted, this time randomly shuffling the decoding target in each repetition, which broke the mapping between the memory load condition and the EEG data.

Finally, we trained separate classifiers on the data of each eccentricity condition. One-way rmANOVAs were used to test for differences between the three eccentricity conditions in decoding performance at each time point (cluster-corrected), the peak performance throughout the retention interval, and its latency.

### Lateralized alpha power

We employed complex Morlet wavelets to analyze the time-frequency decomposition of the epoched EEG signal. Wavelets were generated at 20 evenly spaced frequencies between 6-25 Hz, each with a width of 500 ms. Hence, the number of cycles in each wavelet equaled its frequency divided by 2. We calculated induced power (Kalcher & Pfurtscheller, 1995) by subtracting the averaged evoked response (i.e., the ERP) per experimental condition from the corresponding sing-letrial data, before convolving it with the wavelet, to separate the oscillatory dynamics from phase-locked phenomena. To normalize the single-trial power time courses, the average power during a 200 ms baseline, from −300 to −100 ms relative to cue onset, was subtracted. This early baseline allowed us to avoid that the lateralization of alpha power already in response to the presentation of the arrow cue (i.e., in the time window directly preceding the onset of the memory array; see **Figure 3**) biases the baseline. The lateralized difference in (induced) power per frequency was calculated by subtracting the power time courses of ipsilateral sensors from those on the contralateral side and averaging the results across sensors within the same bilateral ROI used for the CDA analysis, in accordance with the procedure reported by Hakim et al. (2019). Alpha lateralization during vSTM retention has been observed for parietal and occipital channels (e.g., Fukuda et al., 2015; Hakim et al., 2019; Sauseng et al., 2009), suggesting that the ROI used by Hakim et al. (2019) is an appropriate choice of channels to capture such effects. To test the effect of memory load and stimulus eccentricity on alpha power, the results for the frequencies in the alpha range (8–13 Hz) were averaged to form a single time series of lateralized power. We applied the same statistical approach to analyze the effect of the experimental manipulations on this time series as we did for the CDA analysis. The lateralized signal was pooled across all experimental conditions and subjected to *t*-tests at each time point to identify windows in which this signal was different from zero, indicating substantial lateralization. We extracted the averaged signal within the identified clusters (*p* < .05 after cluster-correction for multiple comparisons) and used it as dependent variable for the same statistical test (rmANOVA) as applied to the mean CDA amplitudes.

**Figure 2.**
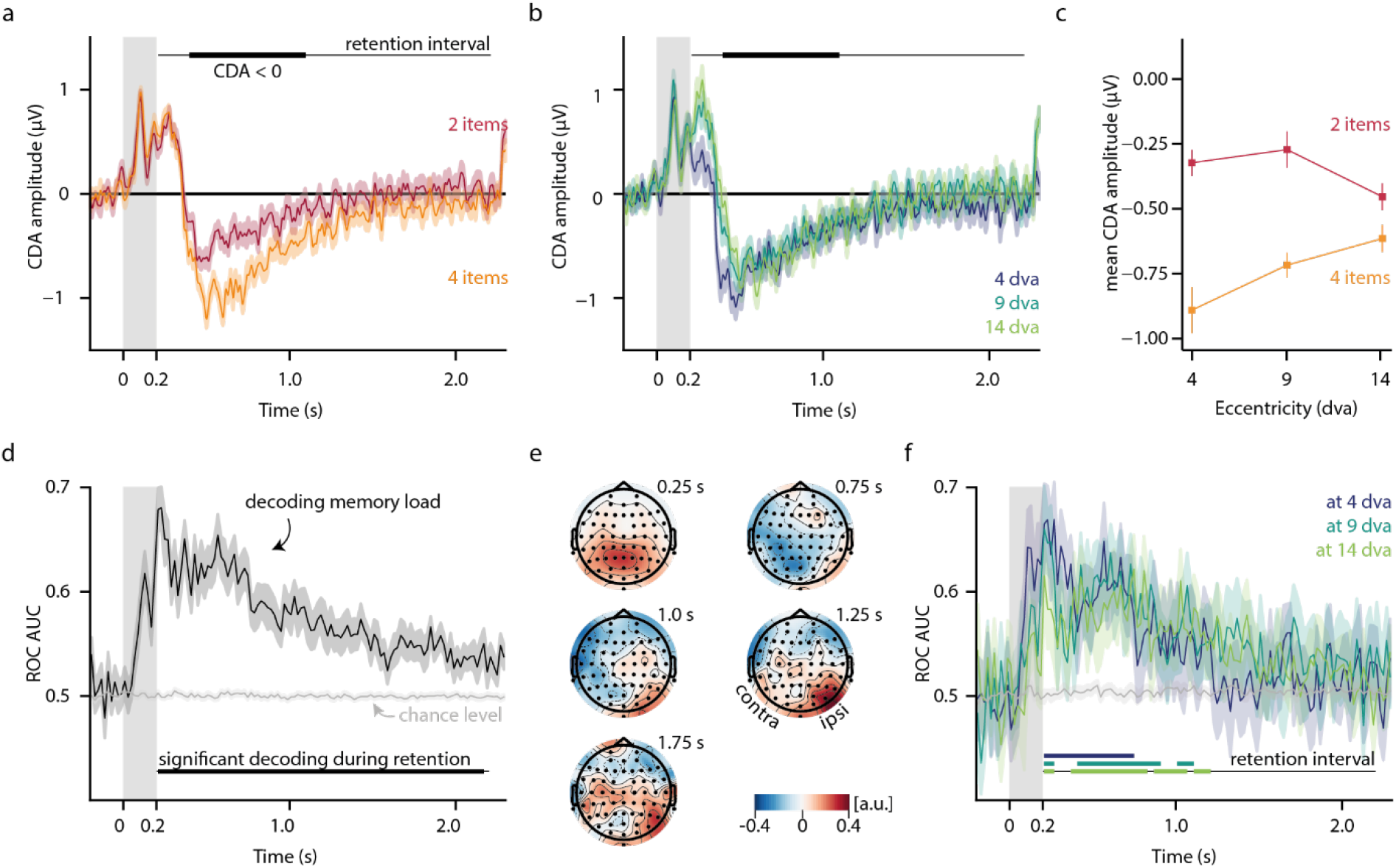
The contralateral delay activity (CDA) plotted for (a) set sizes of two and four memory items and (b) the three stimulus eccentricity conditions. The grey shading indicates the time window during which the stimulus arrays were visible. The thin line at the top of the figure marks the retention interval (i.e., the time window between memory array offset and memory probe onset). The thick line marks the time window of the CDA which was identified using the grand-average across all conditions. (c) The mean CDA amplitudes in this interval per load and eccentricity condition. (d) Time-resolved decoding performance when decoding the memory load from the ERP data. Significance was tested against results from decoding with shuffled labels (light-grey line) and only during the retention interval. (e) The spatial patterns of the classifier for selected time points during the retention interval, normalized and averaged across participants. Since we pooled across the two cue conditions (left vs right hemifield), electrode locations here are relative to the cued side (by convention; left channels: contralateral, right channels: ipsilateral). (f) Time-resolved performance of the classifier decoding memory load separately for the three stimulus eccentricities. The thick colored lines mark the clusters which drove the finding of above-chance decoding. The shaded areas in (a), (b), (d), (f) as well as the error bars in (c) indicate ±1 SEM (taking into account the repeated measures design; Baguley, 2012; Morey, 2008).

**Figure 3.**
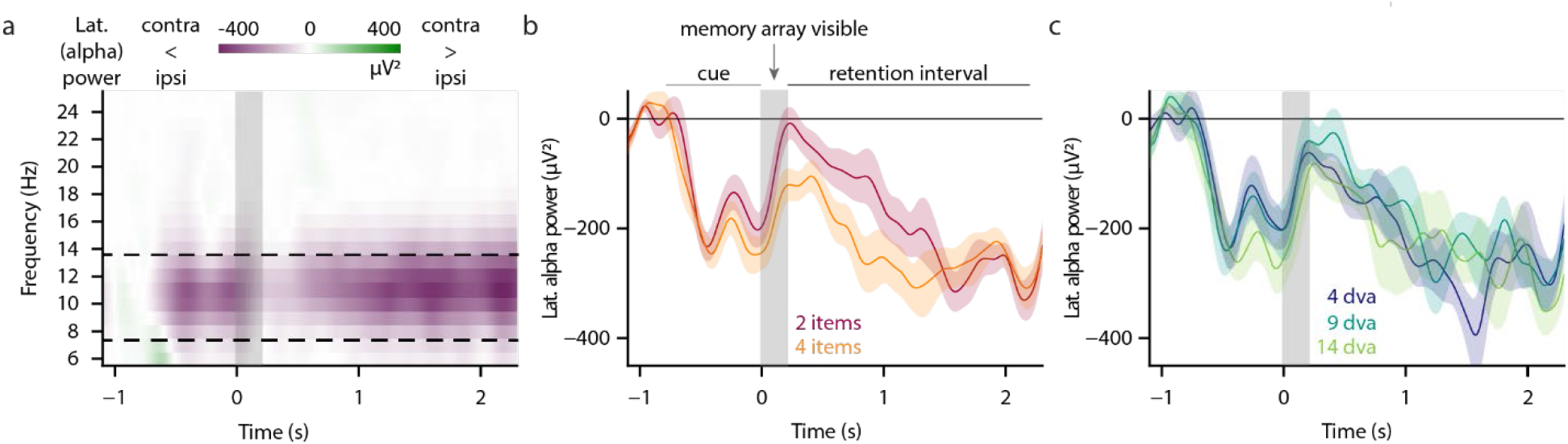
(a) Lateralized power (i.e., after subtracting ipsi-from contralateral induced power time courses, relative to the side of the memory array) for different frequencies. The dashed lines mark the a priori defined alpha frequency range (8–13 Hz). Direction and strength of the lateralization are color-coded. The grey box indicates the time window (0.2 s) during which the memory stimulus was visible. We observed lateralization of power predominantly in the alpha frequency range—both, while the cue was visible (−0.8 to 0 s) and during memory retention (0.2 to 2.2 s). (b, c) The time course of the lateralized power averaged across the alpha frequencies as a function of memory load (b) and eccentricity (c). The shaded areas indicate ±1 SEM (taking into account the repeated measures design; Baguley, 2012; Morey, 2008).

### Decoding from time-frequency data

For further analysis of the time-frequency features of the data, we applied a multivariate decoding approach on the single-participant level, similar to the time-domain analysis. We used the Common Spatial Patterns (CSP) algorithm (Blankertz et al., 2008; Ramoser et al., 2000), as implemented in MNE-Python, which was originally established in the field of brain computer interfaces to extract discriminative spectral features from the EEG for the classification of dichotomous states. CSP solves a Generalized Eigenvalue Decomposition problem by fitting a spatial filter to the EEG data. It allows a linear projection of band-pass filtered data onto components whose power maximally relates to the prevalence of one of two dichotomous states. This allowed us to test the discriminability of trials with low memory load from trials with high memory load by means of their spectral features.

We extracted epochs of 3,000 ms length (−500 to 2,500 ms relative to memory array onset) and subtracted the grand-average ERP from each epoch and channel to remove evoked activity before band-pass filtering (hamming window FIR filter, default settings implemented in MNE) the EEG signals into 10 frequency bands of equal width (2 Hz each), ranging from 6–26 Hz. For each frequency band, we calculated the CSP projection in a coarsely time-resolved fashion by sliding a 500 ms time window in steps of 250 ms (50% overlap) along the epoch. To obtain robust estimates of the covariance matrices necessary for the CSP decomposition, shrinkage regularization (shrinkage parameter *λ* = 0.4) was applied. The logarithmized power of the 6 CSP components with the most extreme eigenvalues formed the feature vector for training a linear discriminant analysis (LDA) classifier to distinguish between trials with low and high memory load. This procedure was embedded in a 5-fold cross-validation to assess the decoding performance. In each fold, the spatial filters for the CSP projection and the LDA weights were fitted to 80% of the data and their discriminability between high and low memory load was tested on the remaining 20% of the trials, using the ROC-AUC as the outcome metric. To yield more robust results, we repeated the cross-validation 10 times, randomly varying the allocation of trials to the training and test set. ROC-AUC values were averaged across folds and repetitions of the cross-validation regime, yielding a time series of ROC-AUC scores for each of the frequency bands.

To assess the chance level for this classification procedure, we calculated the empirical chance performance for each participant by shuffling the decoding targets (memory load condition) at the beginning of each repetition and before applying the same cross-validated decoding pipeline. This yielded a ROC-AUC value for each combination of time window and frequency band when decoding an arbitrary target. For the frequency bands covering the alpha range (8–10, 10–12, and 12–14 Hz) we tested whether the actual decoding performance significantly differed from chance. To this end, we averaged the ROC-AUC time series across these frequency bands and applied, on the group level, cluster-corrected one-sided paired *t*-tests for each time point, following the same principles as for the decoding from the ERP data.

Finally, we compared the three eccentricity conditions by training and testing separate (CSP+LDA) classifiers on the data of each eccentricity condition. To test for significant differences in decoding performance between the conditions, we calculated one-way rmANOVAs per time point (cluster-corrected), for the peak performance in the retention interval, and its latency.

## Results

We tested participants’ memory in a delayed match-to-sample task, manipulating memory load and the eccentricity of the memory array, to assess the neural correlates of visual working memory in a VR setup with concurrent EEG recording. Participants correctly detected a change in *M* = 85.18% of the trials (range: 78.48–92.73%). Memory performance was strongly modulated by memory load and varied only little as a function of eccentricity (**Figure 1b**). Participants showed a significantly higher proportion of correct reports in trials with low (*M* = 94.44%, *SD* = 2.65, 95% CI [92.94, 95.93]), as compared to high memory load (*M* = 75.89%, *SD* = 7.33, 95% CI [74.40, 77.39]; *F*(1,20) = 166.34, *p* < .001). The influence of eccentricity was comparatively small but significant (*F*(2,40) = 4.10, *p* = .024). Participants’ memory performance was lowest for the eccentricity of 14 dva (*M* = 84.00%, *SD* = 5.12, 95% CI [83.23, 84.76]) and significantly worse than memory for stimuli presented at 9 dva (*M* = 85.91%, *SD* = 4.54, 95% CI [85.13, 86.68]) or 4 dva (*M* = 85.68%, *SD* = 4.81, 95% CI [84.98, 86.38]), as corroborated by post-hoc *t*-tests (Δperformance_9–14_ = 1.91%, 95% CI [0.33, 3.49], *t*(20) = 2.52, *p* = .020; Δperformance_4–14_ = 1.68%, 95% CI [0.25, 3.11], *t*(20) = 2.46, *p* = .023). There was no significant difference between the two smaller eccentricities (Δperformance_4–9_ = −0.23%, 95% CI [−1.68, 1.23], *t*(20) = −0.33, *p* = .748). The interaction between memory load and eccentricity did not affect memory performance (*F*(2,40) = 1.45, *p* = .247).

In additional control trials, we assessed whether lower memory performance at the larger eccentricities could be accounted for by diminished stimulus visibility at these eccentricities. Here, the probe appeared immediately after the 200 ms encoding phase (i.e., there was no retention interval). Also in this perceptual control task, the performance was lowest at the largest eccentricity of 14 dva (*M* = 89.46%, *SD* = 5.70, 95% CI [87.82, 91.10]; **Figure 1c**). We corroborated this finding using a two-way rmANOVA and observed a significant main effect of eccentricity (*F*(2,40) = 11.85, *p* < .001). Post-hoc *t*-tests revealed lower performance at 14 dva (*M* = 89.46%, *SD* = 5.70, 95% CI [87.82, 91.10]) as compared to 9 dva (*M* = 94.99%, *SD* = 4.71, 95% CI [93.55, 96.44]; Δperformance_9–14_ = 5.54%, 95% CI [2.33, 8.74], *t*(20) = 3.61, *p* = .002) and 4 dva (*M* = 95.76%, *SD* = 4.23, 95% CI [94.40, 97.13]; Δperformance_4–14_ = 6.30%, 95% CI [3.25, 9.36], *t*(20) = 4.31, *p* < .001). Performance did not differ significantly between the two smaller eccentricities (Δperformance_4–9_ = 0.77%, 95% CI [−1.87, 3.40], *t*(20) = 0.61, *p* = .550). Neither the number of stimuli (low memory load: *M* = 94.13%, *SD* = 4.37, 95% CI [92.81, 95.45]; high memory load: *M* = 92.69%, *SD* = 4.19, 95% CI [91.37, 94.01]; *F*(1,20) = 1.33, *p* = .262) nor the interaction with eccentricity significantly influenced performance in the perceptual task (*F*(2,40) = 1.11, *p* = .338). Thus, although performance in the perceptual task was significantly lower for the largest eccentricity, it was nevertheless very high on an absolute scale. This mirrors the behavioral results in the memory task, where performance was significantly lower for the largest eccentricity, but overall very high with a clear influence of memory load on performance even at the largest eccentricity.

### CDA and PNP

We observed a typical CDA (**Figure 2a**), that is, a lateralization of the ERP which started around 400 ms after onset of the memory array and was enhanced for the higher memory load. Cluster-based permutation testing in combination with the collapsed localizer method (Luck & Gaspelin, 2017) revealed significant lateralization in the a priori defined ROI (electrodes: P3/4, P5/6, PO3/4, PO7/8, O1/2) during the retention interval (200 to 2,200 ms after memory array onset). Two clearly distinguishable clusters drove this effect. Note that the exact temporal extent of these clusters should be interpreted cautiously as cluster-based permutation tests only allow statistical conclusions about the entire population of tested samples (Sassenhagen & Draschkow, 2019). In our analyses, we used the time windows of the clusters as a data-driven proxy to determine intervals which showed a reliably lateralized signal to then assess the influence of memory load and stimulus eccentricity in these time windows. The first of these clusters extended from 200 ms (start of the analyzed time window) to 344 ms after memory array onset (mean amplitude: 0.58 μV, *SD* = 0.56, 95% CI [0.33, 0.84]). The second cluster (mean amplitude: −0.54 μV, *SD* = 0.56, 95% CI [−0.80, −0.29]) extended from 388 to 1,088 ms after stimulus onset. Based on their different polarity and temporal characteristics, we will refer to the earlier cluster as *PNP component* (Papaioannou & Luck, 2020) and to the second one as *CDA component*. The CDA component varied significantly with memory load as corroborated by a two-way rmANOVA (*F*(1,20) = 19.84, *p* < .001). Trials with high memory load evoked significantly larger (i.e., more negative) mean CDA amplitudes (*M* = −0.75, *SD* = 0.66, 95% CI [-1.05, −0.45]) than trials with low memory load (*M* = −0.35, *SD* = 0.53, 95% CI [-0.59, −0.11]). Stimulus eccentricity did not significantly influence the mean CDA amplitude (*F*(2,40) = 0.84, *p* = .439) but interacted with the effect of memory load (*F*(2,40) = 4.55, *p* = .017). Post-hoc paired *t*-tests showed that the difference between trials with low and high memory load was significant for the eccentricities of 4 dva (ΔCDA_low–high_ = 0.57, 95% CI [0.27, 0.87], *t*(20) = 3.97, *p* = .001) and 9 dva (ΔCDA_low–high_ = 0.45, 95% CI [0.21, 0.68], *t*(20) = 3.99, *p* = .001), but not for 14 dva (ΔCDA_low–high_ = 0.16, 95% CI [−0.04, 0.36], *t*(20) = 1.66, *p* = .113). For the mean PNP amplitude, we obtained a different pattern: It was significantly modulated by stimulus eccentricity (*F*(2,40) = 14.93, *p* < .001), with larger eccentricities yielding higher PNP amplitudes. Post-hoc *t*-tests confirmed a significant difference between the mean PNP amplitude at 4 and 9 dva (ΔPNP_4-9_ = −0.41, 95% CI = [−0.62, −0.21], *t*(20) = −4.19, *p* < .001) and between 4 and 14 dva (ΔPNP_4-14_ = −0.53, 95% CI = [−0.78, −0.27], *t*(20) = −4.32, *p* < .001), but not between the two larger eccentricities (ΔPNP_9-14_ = −0.11, 95% CI = [−0.28, 0.05], *t*(20) = −1.44, *p* = .167). Memory load did not have a significant main effect on mean PNP amplitude (*F*(1,20) = 0.01, *p* = .917) nor did it interact with eccentricity (*F*(2,40) = 0.44, *p* = .646).

### Decoding from the ERP

In complementary analyses, we applied a multivariate decoding approach to test whether the memory load (i.e., high vs low) can be decoded from the broadband EEG data during the retention interval in our setup. A cluster-based permutation test, comparing the decoding performance against chance level, yielded a significant difference, driven by a large cluster extending from 209 to 2,169 ms after stimulus onset (**Figure 2d**). We observed the highest decoding performance (ROC-AUC; *M* = 0.79, *SD* = 0.05, 95% CI [0.76, 0.81]) for most participants at *Mdn* = 309 ms (*SD* = 174.31, 95% CI [280.14, 435.00]) after stimulus onset. The topographies of the decoding models (patterns of the classifier normalized per participant and time point, then averaged across participants) confirmed that predominantly parieto-occipital channels were informative for the classification of memory load (**Figure 2e**). The spatial patterns of the linear decoding models varied substantially between participants (see Supplementary Material for individual patterns). Moreover, we determined whether the decoding of the memory load was modulated by eccentricity. To this end, we trained models that predicted memory load separately for each eccentricity. Decoding performance was significantly above chance level during the retention interval for all stimulus eccentricities (**Figure 2f**) and did not vary between the single eccentricities. A cluster-corrected one-way rmANOVA did not reveal any significant differences in memory load decoding performance between the eccentricity conditions at any of the tested time points. Furthermore, there were no significant differences between the eccentricity conditions in peak decoding performance (*F*(2,40) = 2.33, *p* = .110) or the timing of the decoding peak (*F*(2,40) = 1.00, *p* = .375).

### Lateralized alpha power

**Figure 3a** shows the difference in time-frequency decompositions between contra- and ipsilateral regions of interest (ROIs), pooled across cue directions. We observed a significant difference in induced power between contra- and ipsilateral signals during the retention interval, with more power in signals measured from electrodes ipsilateral to the target stimulus. This difference was most prominent in the alpha frequency range and was consistently observed across most participants (see Supplementary Material). As in the ERP analyses, we pooled the data across all experimental conditions (collapsed localizer method; Luck & Gaspelin, 2017) to determine time windows during which the induced power in the alpha range (8–13 Hz) was substantially lateralized. During the retention interval, the lateralization of alpha power was most pronounced for a single cluster extending from 506 ms after memory array onset to the end of the retention interval (i.e., 2,200 ms after memory array onset). We used a two-way rmANOVA to investigate the influence of memory load and stimulus eccentricity on the average lateralization of alpha power (expressed as the difference between contra- and ipsilateral power) in this time window (*M* = −235.36 μV^2^, *SD* = 280.16, 95% CI [−362.89, −107.83]). Neither memory load nor eccentricity significantly influenced the lateralization (memory load: *F*(1,20) = 1.25, *p* = .277; eccentricity: *F*(2,40) = 0.40, *p* = .672; interaction: *F*(2,40) = 0.68, *p* = .515).

### Decoding from time-frequency data

To complement this approach, we applied a multivariate classifier based on CSP transformation and LDA classification to decode memory load from spectral features. More precisely, we trained and tested separate models on a sliding time window during the retention interval and on different frequency bands. The highest decoding performances were observed for the alpha frequency range (8–13 Hz, **Figure 4a**). A cluster-based permutation test confirmed that the decoding was significantly above chance level during the retention interval for these frequencies. This was driven by a single cluster which spanned almost the entire retention interval (**Figure 4b**). However, as mentioned above, the nature of the cluster-based permutation approach and the broad time bins used for feature extraction (500 ms) prohibit a strong interpretation of the start and end times of this cluster as defining the exact time window during which decodable information about memory load is present. Across all participants, the decoding performance (ROC-AUC) was highest from frequencies in the alpha range (*M* = 0.57, *SD* = 0.04, 95% CI [0.55, 0.59]) and in the time bin centered around 1,000 ms (*SD* = 411.88, 95% CI [995.61, 1361.54]) after stimulus onset. Plotting the patterns (i.e., the inverse filter weights; Haufe et al., 2014) of the first CSP component revealed that parieto-occipital channels contributed the most discriminative features, without very clear lateralization (see Supplementary Material for the spatial patterns of the most discriminative CSP component). To investigate the effect of stimulus eccentricity on the success of decoding memory load, we trained and tested separate models on data from the different eccentricity conditions. We found clusters with above-chance decoding performance (*p* < .05) for all three eccentricity conditions (**Figure 4c**). As for the decoding from the broadband ERP data, one-way rmANOVAs contrasting the eccentricity conditions did not indicate significant differences in decoding performance at any time point, in peak performance (*F*(2,40) = 0.19, *p* = .825) or in the timing of the decoding peak (*F*(2,40) = 2.14, *p* = .131).

**Figure 4.**
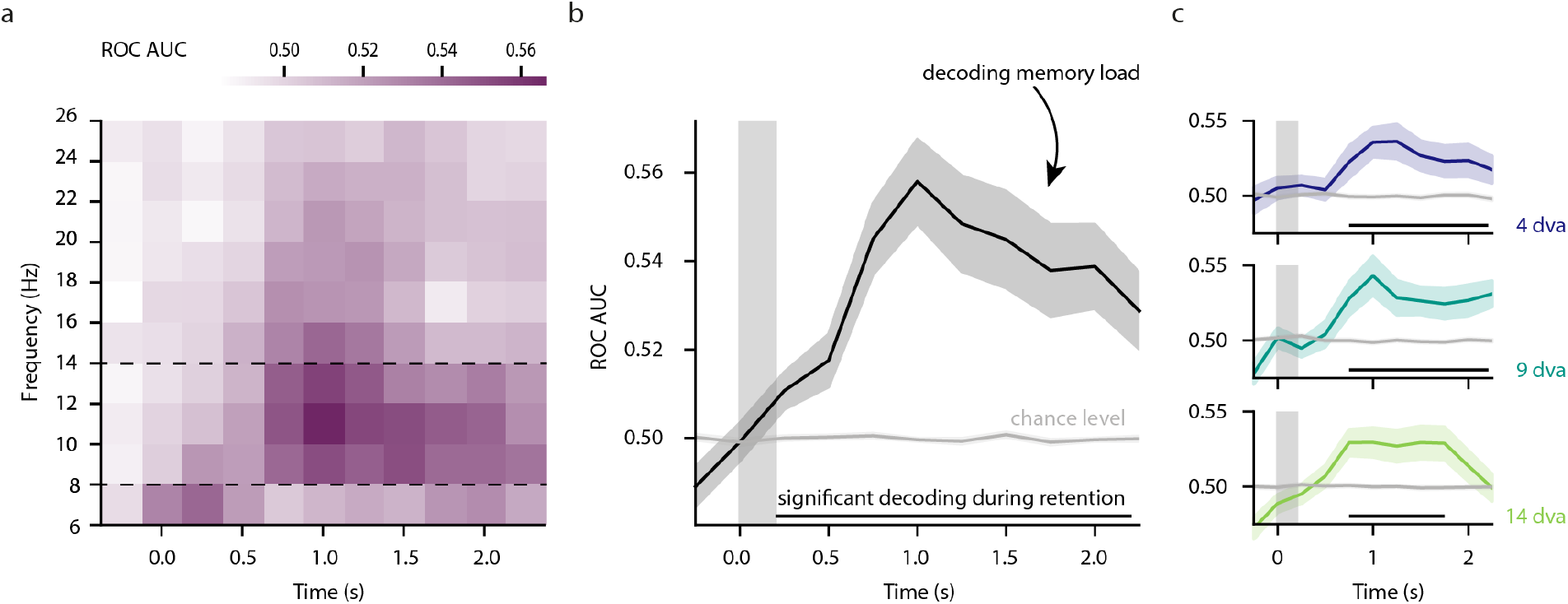
(a) The classifier performance for decoding memory load (CSP+LDA) for different frequency bands and time points. The dotted lines mark the frequency bands that cover the a priori defined alpha frequency range (8–13 Hz). (b) The mean (±1 SEM) decoding performance (classifying memory load) across the frequencies covering the alpha band. Significance was tested against decoding with shuffled labels (grey line) for all time windows covering the retention interval. The horizontal black line at the bottom of the plot indicates the time windows where this difference was significant. (c) Like (b) but decoding memory load separately per stimulus eccentricity condition.

## Discussion

We investigated two well-established EEG components related to visual short-term memory processes, the CDA and lateralized alpha power, in a virtual reality setup. Increasing memory load resulted in diminished memory performance and we observed both a pronounced CDA and a lateralization of alpha power during the retention interval. Moreover, we explored potential spatial constraints by varying the horizontal stimulus eccentricity which played a distinct role for these correlates of visual short-term memory. While we observed the CDA as well as a lateralization of alpha power during memory retention at all tested eccentricities, the characteristic memory load effect on the CDA amplitude was present only for the two smaller eccentricities (4 and 9 dva) but not for stimuli with an eccentricity of 14 dva. To corroborate these findings, we used multivariate approaches to decode the memory load from the EEG signals. This analysis revealed information about the memory load across all tested eccentricities, both when decoding from broadband ERP data and from alpha power signals.

Participants’ memory performance decreased for trials with higher memory load, and was similar as in previous (non-VR based) studies that tested comparable memory loads (e.g., Feldmann-Wüstefeld, 2021). Memory performance decreased for stimulus arrays at an eccentricity of 14 dva as compared to the smaller eccentricities (4 and 9 dva), and the influence of memory load on performance was comparable at all three eccentricities. Importantly, performance in the perceptual task mirrored the role of eccentricity in the memory task, suggesting that the decline in memory performance at an eccentricity of 14 dva can be accounted for, at least in part, by perceptual rather than memory-related processes. Encoding difficulties might arise from the VR headset (e.g., the Fresnel lens setup in the HTC Vive Pro Eye makes the perceived image substantially blurrier towards the edges). In preliminary tests of the setup, we observed that stimuli presented at eccentricities beyond 15 dva were increasingly blurred and harder to perceive. Therefore, we kept the center of the stimulus arrays at 14 dva or smaller. However, since the stimuli were spatially jittered around the array centers, single stimuli in the 14 dva condition might have appeared blurry—possibly resulting in worse perception in such trials. Alternatively, the lower perceptual performance at the largest eccentricity could simply reflect coarser resolution of human visual perception toward the periphery of the visual field (Anstis, 1998) where the receptive fields of neurons are larger than in the fovea and parafovea (Freeman & Simoncelli, 2011). Also the perception of color declines in a similar manner towards the periphery of the visual field (Hansen et al., 2009; Noorlander et al., 1983; Tyler, 2015). Increasing the stimulus size as a function of eccentricity, to counteract the cortical magnification (Cowey & Rolls, 1974), might improve performance at the largest eccentricity. However, in most VR paradigms (as well as in everyday life), the size of a stimulus is a constant and not dependent on its position relative to the observer. One of the goals of our study was to provide parameters for such, more naturalistic, VR experiments which is why we decided against scaling the stimuli as a function of eccentricity. Overall, the behavioral results demonstrate that participants were able to perform the task with a VR headset and that the manipulation of memory load yielded the intended effect at all eccentricities.

In studying the neurophysiological concomitants of visual short-term memory, we found a clear CDA and its expected amplitude scaling with memory load, that is, more negative amplitudes in trials with higher memory load. Therefore, the interferences with the EEG signals caused by wearing a VR headset are not an insurmountable obstacle for studying the CDA in immersive VR setups. In comparison to other VR-based paradigms, our study was still highly controlled in the sense that participants did not move during the performance of the task and that the stimuli were minimalistic. In settings that require more movements or use more complex stimuli—which is the case for many VR paradigms—noise levels may be substantially higher and the CDA harder to find. While other EEG components have been observed successfully in experiments where participants could move more freely than in our study (e.g., Hofmann, Klotzsche, Mariola et al., 2021; Krugliak & Clarke, 2022; Liang et al., 2018; Tauscher et al., 2019), it will be important to determine whether also the CDA can be observed in less constrained VR setups. Especially in settings for which one expects higher levels of physiological and movement-related noise (i.e., a lower signal-to-noise ratio), one should aim for a sufficiently large sample size or number of trials to allow detecting the CDA or an influence of memory load on the CDA (see Ngiam et al., 2020 for a helpful guideline).

Stimulus eccentricity played a critical role for the influence of memory load on the CDA amplitude. More specifically, we did not observe a significantly different CDA amplitude between the two memory loads at the largest eccentricity (14 dva). Our results do not allow concluding whether this negative finding resulted from a reduced CDA in the high load condition, a stronger CDA in the low load condition, or both, as the post-hoc tests remained inconclusive in this regard. Perceptual effects—that might have led to the small decrease in behavioral performance for the largest eccentricity—could also underlie the effect on the CDA. This effect is particularly relevant for the condition with a greater number of stimuli, since here more stimuli are positioned in a perceptually problematic eccentricity range. Therefore, in some trials of the high memory load condition, participants might have encoded and maintained only a fraction of the displayed stimuli, which in turn would be reflected in a similar CDA as in the low memory load condition. Another interpretation is that hampered perception (e.g., of colors) in the periphery leads to altered ERPs and that such an effect is more pronounced for conditions with a higher number of stimuli. However, Papaioannou & Luck, (2020) found that scaling their stimuli according to cortical magnification did not affect lateralized EEG components, and there was no interaction with the effect of stimulus eccentricity in particular. Another possible explanation is that the CDA was—at least partially—overlaid by the PNP component, which exhibits an opposite polarity. Indeed, we found that the amplitude of the PNP increases with larger stimulus eccentricities, therefore replicating the finding of Papaioannou & Luck (2020). Given that the two components are superimposed in sensor space, higher stimulus eccentricity might lead to a weaker CDA. Trials with more stimuli (i.e., higher memory loads) may be affected disproportionally stronger. In an exploratory analysis (see Supplementary Material) we revealed that stimulus eccentricity particularly affected the early parts of the CDA time window, where also the strongest load-effects on the CDA are observed. This is relevant for studies in which eccentricity of the stimuli varies between trials: Stimulus eccentricity will influence early, attention related components (e.g., the PNP) which in turn might bias the CDA results.

There was a robust lateralization of alpha power during the retention interval, with less power in contralateral than in ipsilateral signals. However, we did not find any effect of memory load on alpha lateralization, contrary to previous research (Sauseng et al., 2009; but see also Figueira et al., 2020; Fukuda et al., 2016). This discrepancy may be due to a low signal-to-noise ratio for alpha power signals resulting from our VR-EEG setup or the manipulation of stimulus eccentricity. The clearly accentuated alpha peak in the power spectrum observed for all eccentricity conditions (see Supplementary Material) speaks against this interpretation but cannot fully refute it. Alternatively, insufficient power in the study design may have contributed to the absence of an effect of memory load on lateralized alpha power. However, our study included a comparable number of participants and a substantially larger number of trials than the study of Sauseng et al. (2009) who reported such effects. Ultimately, the lateralization of alpha power in (lateralized) visual short-term memory tasks might reflect the unilateral allocation of attention rather than memory processes (Fukuda et al., 2015; Hakim et al., 2019; Wang et al., 2019) and therefore be independent of the memory load. The link between the spatial focus of attention and the pattern of alpha power measured with EEG is well-established (Foster et al., 2017; Worden et al., 2000). Our findings align with the idea that the lateralization of alpha power during the retention interval reflects a shift of attention, as evidenced by the similar pattern of alpha lateralization in response to the lateralized cue, observed before any memory processes were required (**Figure 3**). Furthermore, the lateralization of alpha power increased towards the end of the retention interval (while the memory load-dependent CDA peaked substantially earlier), possibly because participants shifted their attention to the expected location of the memory probe. Similar shifts of attention have been observed when participants prepare saccades to a lateral, parafoveal targets (for example during left-to-right reading; Kornrumpf et al., 2017). While we cannot draw definite conclusions about the relationship between attention, memory processes, the lateralization of alpha power, and the CDA, our experiment shows that VR technology can be used to study them in novel and more complex experimental settings. Combining the existing knowledge about the CDA and alpha lateralization with the new capabilities of VR headsets, such as large visual fields, stereoscopic stimulation or the integration of head and body movements in experimental designs, will yield new insights into the characteristics and interrelation of these components.

We complemented our univariate analyses with multivariate approaches, by decoding memory load from EEG signals during the retention interval. These methods have been shown to be more sensitive than univariate approaches, as they combine EEG channels in an optimal way to solve classification or regression problems (Adam et al., 2020). Above-chance decoding performance indicates—in a time-resolved fashion—that the EEG contains information about the decoding target. We were able to decode the memory load from both, the phase-locked data (i.e., the ERP) as well as from induced alpha power (i.e., the result of the time-frequency decomposition of the signal) throughout the retention interval. Compared to our univariate analyses of lateralized signals (i.e., hemispheric differences) in a pre-defined ROI, the multivariate classifiers were trained on data from all channels, allowing them to pick up non-lateralized features and information from channels outside the ROI. The complementary nature of this approach became particularly evident when investigating effects on alpha power, where we did not observe a significant effect of memory load in the univariate analysis but were able to decode the level of memory load using the multivariate classifier. Furthermore, the decoding performance and lateralized alpha power showed different time courses— the decoding performance peaked in the middle of the retention interval, while the lateralization of alpha power increased towards the end. This suggests that the decoding exploits additional, likely non-lateralized, features. Interestingly, the decoding performance was independent of stimulus eccentricity for both the time-locked signals (ERP) and the time courses of alpha power, despite being a more sensitive measure than univariate approaches. Therefore, the perceptual confounds that slightly impacted behavioral memory performance at the largest eccentricity may not fully explain the interaction between stimulus eccentricity and memory load in the CDA analysis. If the largest eccentricity systematically led to the encoding and maintenance of less information during the retention interval, particularly for trials with high memory load, the classifier should have performed worse on data from these trials. Overall, our decoding results show that the ERP and induced alpha power contain information about memory load during the retention interval and across all tested eccentricities. Multivariate approaches can be effective at detecting this information, particularly in conditions of low signal-to-noise ratios.

One of the benefits of using VR setups for experimentation is the ability to obtain and utilize a larger number of measures in comparison to many classical lab setups (Draschkow, 2022). VR headsets, for example, allow for real-time tracking of head position and orientation. We used this feature to ensure a reliable craniotopic presentation of our stimuli. To achieve this (i.e., a stable spatial relation between stimulus and observer) in conventional lab setups, experimenters typically need to constrain participants’ head movements (for example, by using a chin rest) which can be inconvenient and unnatural. In a VR setup, experimenters can easily incorporate head movements instead of physically restricting them. Furthermore, there is a growing number of VR hardware products with built-in eye tracking capabilities. We used the eye tracker in our VR headset to ensure (post-hoc) that participants maintained stable fixation and to exclude trials where this was not the case. In future studies, VR headsets with eye tracking capabilities could be used for gaze-contingent paradigms, presenting stimuli in relation to both the head position and the current direction of the eye gaze. However, currently available VR hardware and software solutions still pose challenges (e.g., high latencies) for adapting these techniques to scientific standards (Stein et al., 2021).

VR is increasingly being used to study visual short-term memory (e.g., Draschkow et al., 2020, 2022; Thom et al., 2022) by offering a practical and well-controlled way to conduct experiments which incorporate (some of) the complexities of everyday life and to use its sophisticated tracking and stimulation opportunities. Up to now, most of these studies focus on behavioral and eye tracking measures. Our findings encourage the use of VR to measure EEG signals related to visual short-term memory and attention (e.g., the CDA or the lateralization of alpha power), and identify caveats for future experimental designs.

## CRediT statement

**Felix Klotzsche:** Conceptualization, Data curation, Formal Analysis, Investigation, Methodology, Software, Visualization, Writing – original draft, Writing – review & editing;

**Michael Gaebler:** Funding Acquisition, Resources, Supervision, Writing – review & editing;

**Arno Villringer:** Resources, Writing – review & editing;

**Werner Sommer:** Methodology, Supervision, Writing – review & editing;

**Vadim Nikulin:** Methodology, Supervision, Writing – review & editing;

**Sven Ohl:** Conceptualization, Formal Analysis, Visualization, Writing – original draft, Writing – review & editing;

## Acknowledgments

We thank Zoya Mooraj and Sean Larpoon for their help with data acquisition, Jeroen de Mooij for valuable input regarding the implementation of the experiment in Unity3D, and ChatGPT for support with code documentation and revising parts of the manuscript. This research was supported by a DFG research grant to S.O. (OH 274/2-2), the German Federal Ministry for Education and Research (13GW0206), and the cooperation between the Max Planck Society and the Fraunhofer Gesellschaft (grant: project NEUROHUM). The authors declare no competing financial interests.

