## Supplementary Material for "Visual short-term memory related EEG components in a virtual reality setup"

### Control analyses

To exclude the possibility that the effects of memory load and eccentricity on the CDA were specific to our selection of the CDA time window (which we gained from the data-driven temporal localizer approach), we conducted a set of supplementary analyses. First, we calculated the mean CDA using the time window reported by Hakim et al. (2019). This alternative time window (400 to 1,450 ms after stimulus onset) started (slightly) later and lasted (substantially) longer than the one yielded by the temporal localizer approach (388 to 1,088 ms). In line with our results, we observed also for this alternative time window that the CDA mean amplitude varied significantly with memory load ($F(1,20) = 14.74$, $p = .001$) while stimulus eccentricity did not significantly influence mean CDA amplitude ($F(2,40) = 0.45$, $p = .638$). In contrast to the analysis based on the data-driven time window, there was no significant interaction of eccentricity with the effect of memory load ($F(2,40) = 3.04$, $p = .059$). Post-hoc paired *t*-tests, however, revealed the same pattern as found in the main analysis: the difference between trials with low and high memory load was significant for the eccentricities of 4 dva ($\Delta CDA_{low–high} = 0.47$, 95% CI [0.16, 0.78], $p = .004$) as well as 9 dva ($\Delta CDA_{low–high} = 0.39$, 95% CI [0.16, 0.62], $p = .002$), but not for 14 dva ($\Delta CDA_{low–high} = 0.14$, 95% CI [–0.06, 0.33], $p = .155$). To further explore this pattern with a more sensitive analysis approach, we calculated a hierarchical linear mixed-model (using the statistical software package *lme4*; Bates et al., 2015) which takes single-trial data into account and allows for modeling random intercepts per participant. We modeled the mean CDA amplitude (averaged across channels in our ROI and time points of the respective time window) as a function of memory load (treatment contrast; set size of 2 items as a baseline), eccentricity (treatment contrast; 9 dva as a baseline) as well as their interaction. We fitted one of these models separately for the mean CDA calculated for the time window identified with the temporal localizer method and for the alternative time window, and we determined the significance of the model predictors using the R package lmerTest (Kuznetsova et al., 2017).

This analysis revealed the same pattern of results for both time windows (Table S1): memory load had a strong influence on the mean CDA amplitude while eccentricity did not have a significant main effect. The interaction effect between memory load and eccentricity was small (compared to the main effect of memory load) but significant for both time windows and was driven by the trials with the largest stimulus eccentricity (14 dva) which exhibited a smaller difference between the memory load conditions as compared to trials with 9 dva or 4 dva. There was no significant difference between the two smaller eccentricity conditions. This is the same pattern that we observed with the (non-hierarchical) rmANOVA paired with post-hoc *t*-tests for the data-driven time window. We suspect that we did not observe a significant interaction effect of memory load and eccentricity with the rmANOVA for the a priori time window due to a lack of statistical power. We will discuss in the following paragraph why the interaction effect is smaller in the alternative time window. The more sensitive mixed-model approach allows to detect also weaker effects. Overall, we conclude that the results gained with the original time window (in accordance with the mixed-model) represent a useful while parsimonious description of the data. The key insight: for the largest eccentricity (14 dva), the load effect on the CDA was substantially weakened as compared to the two smaller eccentricities (Figure S1a).

Based on Figure S1a, we assumed that the interaction effect was weaker for the alternative time window as eccentricity might predominantly affect early components of the ERP (like the PNP component). To address this concern, we conducted a time-resolved version of the rmANOVA approach by fitting the according model ($CDA \sim MemoryLoad * Eccentricity$) separately for each sample during the retention interval (Fig. S1b). This revealed different time courses for the main effects of memory load and eccentricity, with the latter peaking substantially earlier, before the onset of the investigated CDA time windows. As expected, the interaction effect only plays a role during the early phases of the CDA time windows and wears of quickly. By using a longer (and/or later) time window to calculate the mean CDA amplitude, the proportion of samples affected by the interaction effect decreases. As the main effect of memory load is stronger and more long lasting, it can also be found more easily in such longer time windows. The post-hoc $t$-tests, however, did not show differences in CDA mean amplitude for either of the time windows, suggesting that increasing the length of the interval is not sufficient to counteract the effect of the interaction.

### Supplementary figures & tables

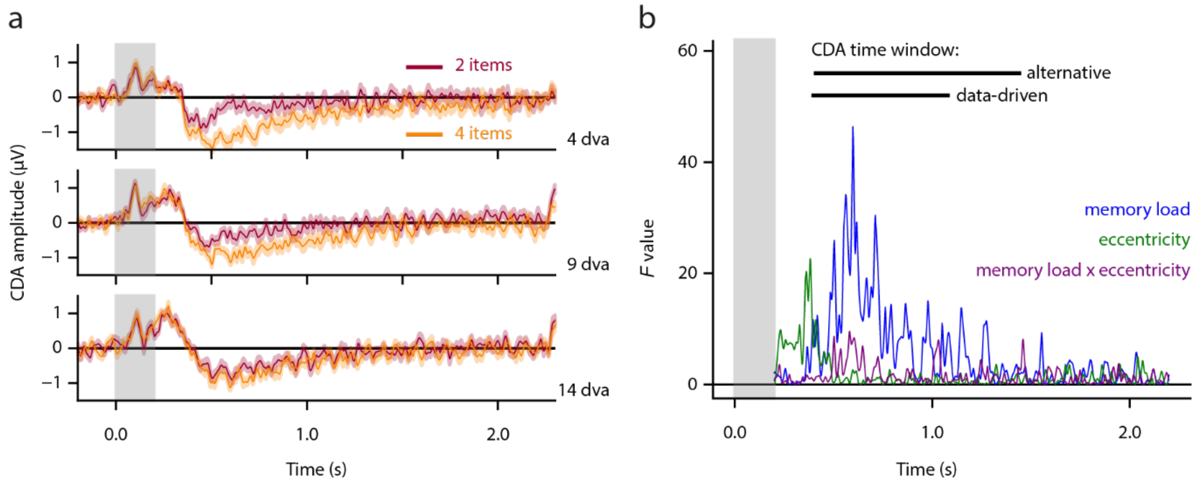


***Figure S1:*** (a) The effect of memory load (2 vs 4 items) on the CDA per eccentricity level. (b) Effects of a time-resolved rmANOVA, modelling the CDA (i.e., lateralized ERP in our ROI) as a function of memory load, eccentricity, and their interaction. It is important to stress the exploratory nature of this supplementary analysis. Therefore, we refrained from performing significance tests.

**Table S1**

*Effects in the mixed-model analysis of the mean CDA amplitude for two different time windows*

| | predictor | b | *t* | df | 95% CI |
|---|---|---|---|---|---|
| CDA time window: 388 – 1088ms | Intercept (2 items; 9 dva) | –0.26 | –1.87 | 34.81 | [–0.55; 0.02] |
| | Memory load (4 vs 2 items) | –0.50 | –4.73*** | 13358.30 | [–0.71; –0.29] |
| | Eccentricity (4 vs 9 dva) | –0.05 | –0.45 | 13358.48 | [–0.26; 0.16] |
| | Eccentricity (14 vs 9 dva) | –0.19 | –1.77 | 13358.38 | [–0.39; 0.02] |
| | Memory load x Eccentricity (4 vs 2 items) x (4 vs 9 dva) | –0.08 | –0.54 | 13358.48 | [–0.38; 0.22] |
| | Memory load x Eccentricity (4 vs 2 items) x (14 vs 9 dva) | 0.35 | 2.37* | 13358.37 | [0.06; 0.65] |
| CDA time window: 400 – 1450ms | Intercept (2 items; 9 dva) | –0.17 | –1.28 | 38.38 | [–0.44; 0.10] |
| | Memory load (4 vs 2 items) | –0.44 | –4.08*** | 13358.33 | [–0.66; –0.23] |
| | Eccentricity (4 vs 9 dva) | –0.04 | –0.36 | 13358.55 | [–0.25; 0.17] |
| | Eccentricity (14 vs 9 dva) | –0.19 | –1.72 | 13358.42 | [–0.40; 0.03] |
| | Memory load x Eccentricity (4 vs 2 items) x (4 vs 9 dva) | –0.04 | –0.26 | 13358.55 | [–0.34; 0.26] |
| | Memory load x Eccentricity (4 vs 2 items) x (14 vs 9 dva) | 0.32 | 2.12* | 13358.42 | [0.02; 0.62] |

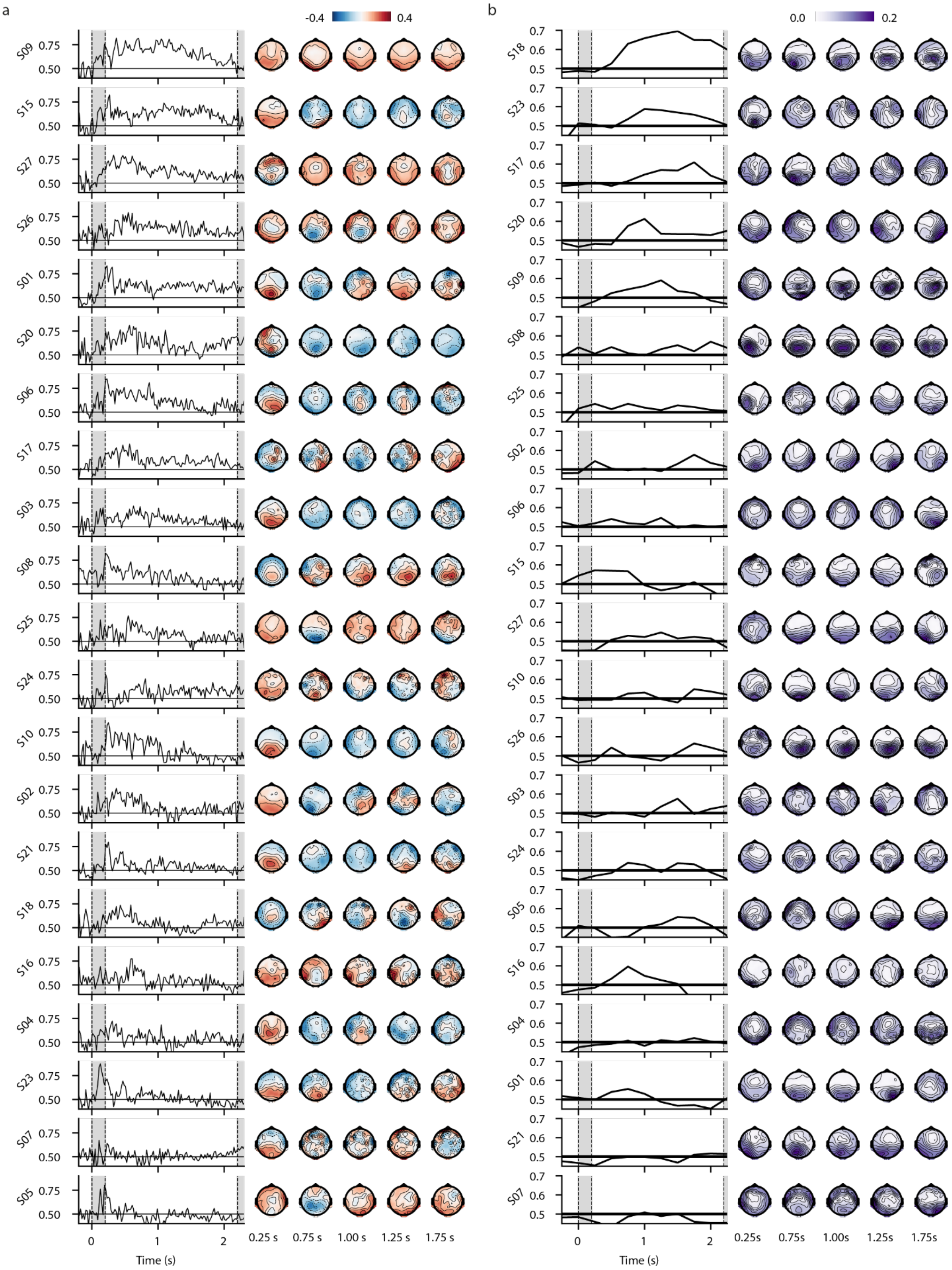

a
-0.4
0.4
S09
S15
S27
S26
S01
S20
S06
S17
S03
S08
S25
S24
S10
S02
S21
S18
S16
S04
S23
S07
S05
0.75
0.50
0
1
2
Time (s)
0.25 s
0.75 s
1.00 s
1.25 s
1.75 s
b
0.0
0.2
S18
S23
S17
S20
S09
S08
S25
S02
S06
S15
S27
S10
S26
S03
S24
S05
S16
S04
S01
S21
S07
0.7
0.6
0.5
0
1
2
Time (s)
0.25s
0.75s
1.00s
1.25s
1.75s

***Figure S2:*** Time courses of the decoding performance and the associated spatial patterns for the single participants. Participants are sorted, separately for each subfigure, decreasingly by the maximum average decoding performance in the time window in which the overall decoding performance (across participants) was significantly above chance level. (a) Decoding from the broadband EEG data (ERP). (b) Decoding from time-frequency data (here: alpha frequencies 8–14 Hz). As we analyzed induced power, which reflects the non-phase locked signal from an oscillating dipole (i.e., with arbitrary polarity at a given point in time), we show the absolute pattern weights to avoid cancellation across participants and repetitions.

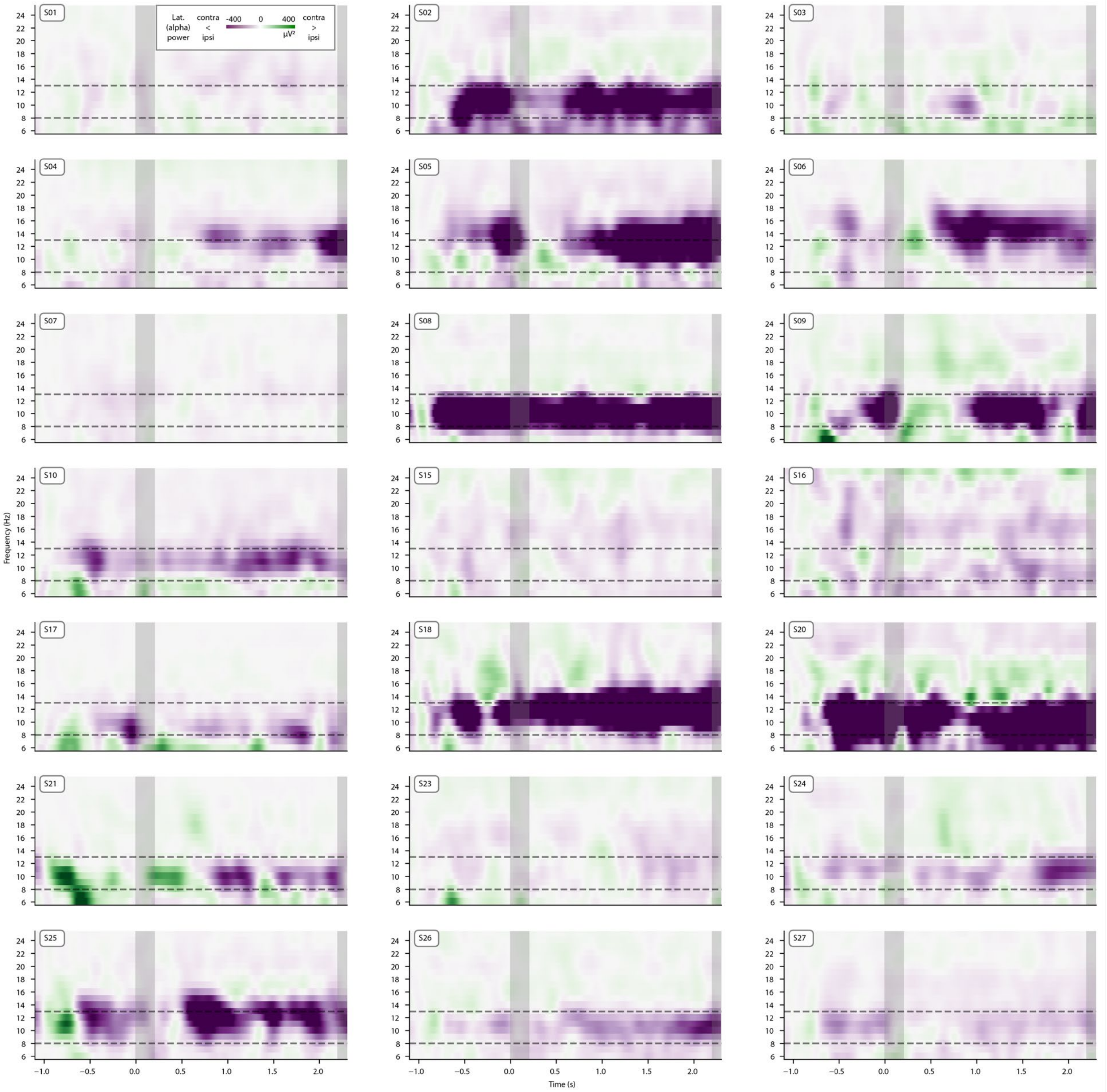


*Figure S3:* Lateralized time-frequency data (difference between contra- and ipsilateral channels) for each participant. The dotted lines mark the lower and upper limit for the frequency range that we included in our analyses regarding alpha lateralization.

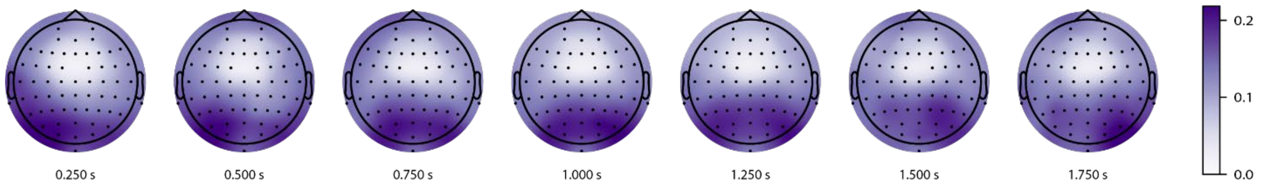


***Figure S4:*** Averaged absolute pattern weights (gained by multiplying the covariance of the EEG data with the filter weights; Haufe et al., 2014) of the most discriminative CSP component (normalized per time-bin and participant before averaging) for the alpha range. As we analyzed induced power, which reflects the non-phase locked signal from an oscillating dipole (i.e., with arbitrary polarity at a given point in time), we show the absolute pattern weights to avoid cancellation across participants and repetitions.

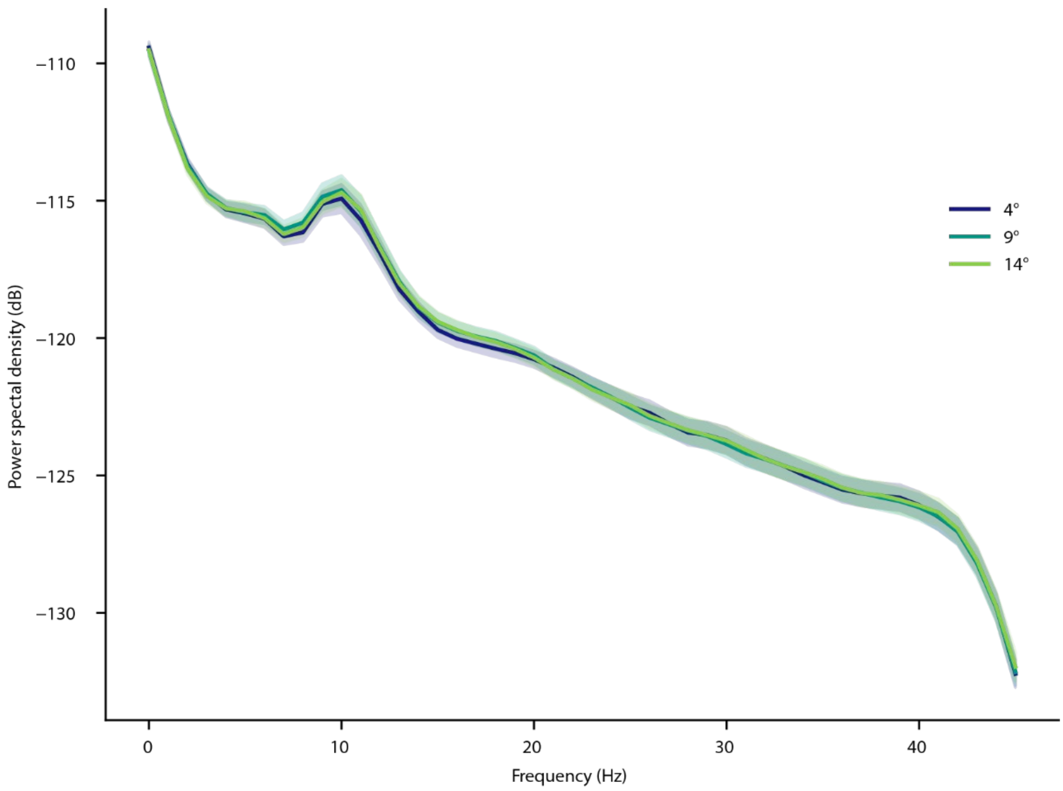


***Figure S5:*** Power spectral density (PSD) per eccentricity condition during the retention interval (shaded area: standard error). Data stems from the bilateral channel-pairs which formed the ROI for the calculation of the CDA and lateralized alpha power. We used Welch's method to calculate the PSD with a window size of 512 samples. An alpha peak is recognizable for each of the eccentricity conditions with no evident difference between the conditions.
